# Sharing of heteroplasmies between human liver lobes varies across the mtDNA genome

**DOI:** 10.1101/155796

**Authors:** Alexander Hübner, Manja Wachsmuth, Roland Schröder, Mingkun Li, Anna Maria Eis-Hübinger, Burkhard Madea, Mark Stoneking

## Abstract

Mitochondrial DNA (mtDNA) heteroplasmy (intra-individual variation) varies among different human tissues and increases with age, suggesting that the majority of mtDNA heteroplasmies are acquired, rather than inherited. However, the extent to which heteroplasmic sites are shared across a tissue remains an open question. We therefore investigated heteroplasmy in two liver samples (one from each primary lobe) from 83 Europeans, sampled at autopsy. Minor allele frequencies (MAF) at heteroplasmic sites were significantly correlated between the two liver samples from an individual, with significantly more sharing of heteroplasmic sites in the control region than in the coding region. We show that this increased sharing for the control region cannot be explained by recent mutations at just a few specific heteroplasmic sites or by the possible presence of 7S DNA. Moreover, we carried out simulations to show that there is significantly more sharing than would be predicted from random genetic drift from a common progenitor cell. We also observe a significant excess of non-synonymous vs. synonymous heteroplasmies in the coding region, but significantly more sharing of synonymous heteroplasmies. These contrasting patterns for the control vs. the coding region, and for non-synonymous vs. synonymous heteroplasmies, suggest that selection plays a role in heteroplasmy sharing.

## INTRODUCTION

The mitochondrial genome is present in many copies in a single cell, and inter-individual variation in the mitochondrial genome of an individual is called mtDNA heteroplasmy (Larsson 2010). In humans, it has been shown that detrimental mtDNA mutations are usually present in a heteroplasmic state at low frequencies, with high frequencies of the deleterious allele leading to functional defects and a disease phenotype (Larsson 2010; Wallace and Chalkia 2013; Stewart and Chinnery 2015). In addition, heteroplasmy is a general phenomenon in aging individuals, where the minor allele is present at rather low frequencies (often below 4%) and many of the affected sites are part of the control region (Stewart and Chinnery 2015). The total number of heteroplasmic sites strongly correlates with age and several studies have shown that heteroplasmic sites are tissue specific (Michikawa et al. 1999; Calloway et al. 2000; Wang et al. 2001; Samuels et al. 2013; Li et al. 2015; Naue et al. 2015), i.e. sites which are frequently heteroplasmic in one tissue are homoplasmic in all other tissues of the same individual.

The tissue specificity of heteroplasmic sites and the association between the number of heteroplasmies and age would suggest that the majority of heteroplasmies are not inherited from the previous generation but are acquired during the lifetime of an individual in a tissue-dependent manner. Therefore, the question arises, whether cells from the same tissue share a similar profile of heteroplasmic sites despite being separated for many cell divisions? Studies on mouse embryos containing two different mtDNA haplotypes have shown that mtDNA segregation occurs rapidly between generations and the distribution of mtDNA haplotypes in the F1 generation resembles a pattern expected under random genetic drift (Jenuth et al. 1996). This result is expected when the underlying heteroplasmies are evolving neutrally. However, it has been shown that mtDNA variants do not behave fully neutrally (Nachman et al. 1996; Jenuth et al. 1997) and more recent studies on human cells indicated that the observed variance in heteroplasmic levels between cells is less stochastic than expected by random genetic drift (Raap et al. 2012; Jayaprakash et al. 2015). This suggests that there might be further population genetic forces, e.g. within-cell mtDNA population structure (Kowald and Kirkwood 2011) or selection, that shape the distribution of heteroplasmies within as well as between tissues.

While most age-related heteroplasmies occur in the control region, human liver tissue is unusual in showing an excess of heteroplasmies involving non-synonymous mutations in the mtDNA protein-coding genes (Li et al. 2015). This result is remarkable because these coding region mutations are likely to have a functional effect (Li et al. 2015), and coding region mutations are strongly selected against during transmission from one generation to another (Stewart et al. 2008). Thus, it seems that mtDNA in human liver tissue exhibits a relaxation of purifying selection and age-related positive selection for somatic mutations that decrease mitochondrial function (Li et al. 2015). This makes liver a good candidate tissue to analyze the sharing of heteroplasmic sites with respect to different evolutionary forces.

While some studies have investigated the amount of variation in levels of heteroplasmy in cells that arose from a single ancestor cell in cell culture (Raap et al. 2012; Jayaprakash et al. 2015), to date there has been no such investigation comparing heteroplasmy across a tissue. We therefore obtained one blood sample and two liver samples (one from each primary lobe) from 83 Europeans, sampled at autopsy. MtDNA heteroplasmy was evaluated by capture-enrichment sequencing (Li et al. 2010; Maricic et al. 2010; Li and Stoneking 2012; Li et al. 2015), and we analyzed sharing of mtDNA heteroplasmy between the liver lobes for different regions of the mitochondrial genome. We find a high correlation in the minor allele frequency (MAF) at heteroplasmic sites in the control region between the two liver samples, but a much weaker correlation in the coding region.

## RESULTS

### Heteroplasmy sharing within the liver

We investigated mitochondrial DNA heteroplasmy in liver and blood tissue samples of 83 individuals. For liver, two samples taken from different lobes were analyzed in order to compare the heteroplasmic pattern in different parts of the tissue. The samples were capture-enriched for mtDNA and sequenced to an average sequencing depth of 1,175-fold for blood samples and 2,640-fold for liver samples. Applying a threshold of 2.5 % MAF, we detected 541 heteroplasmic sites at 343 different positions (Supplementary Table S1). More heteroplasmic sites were observed in the coding region for liver (280 sites) compared to blood (64 sites), but the most abundant heteroplasmic sites in liver were in the control region (site 72: 67 individuals, site 60: 26 individuals, site 94: 20 individuals), which were only rarely observed in blood (site 72: 1 individual, site 60: 1 individual, site 94: no individuals). These data are in accordance with results from a previous study ((Li et al. 2015), Supplementary Figure S1), indicating that heteroplasmy is tissue specific, with different individuals exhibiting similar heteroplasmic patterns.

Virological tests revealed that three individuals had active hepatitis B virus infection, one had active hepatitis C virus infection and one individual was HIV positive, with low viral load (Supplementary Table S2). Those individuals were kept in all downstream analyses, as the number of positive cases was too low to analyze separately. There was no effect of liver fat content on either the total number or the MAFs of heteroplasmic sites (Supplementary Figure S2, p>0.05) and the mitochondrial DNA copy numbers, estimated for each liver sample as described before (Wachsmuth et al. 2016), were highly correlated between corresponding liver samples of an individual, suggesting no functional differences between the liver lobes (Supplementary Figure S3, r=0.81, p<0.001).

We next asked whether heteroplasmy was correlated between the three different samples from an individual. For blood and each liver sample, there is a low but significant correlation (r=0.35 and 0.38, p<0.001, Supplementary Figure S4) with only 7.5% of the heteroplasmies shared between the tissues. However, the correlation between the two liver samples was higher (Figure 1A, r=0.90, p<0.001). While 355 sites were heteroplasmic in only one of the two liver samples of an individual, 136 sites (28%) were heteroplasmic in both liver samples and these exhibited similar MAFs (Figure 1A). Moreover, there were more shared heteroplasmies from the control region than from the rest of the genome (Table 1, p<0.001): 55% of control region heteroplasmies were shared, vs. 7.5% of heteroplasmies outside the control region. The high correlation between the MAFs in the two liver samples was mainly driven by the control region (Figure 1B, r=0.90, p<0.001), while there was a lower, but still significant correlation outside the control region (Figure 1C, r=0.29, p<0.001). The difference in correlation coefficients is significant, based on random partitions of all of the heteroplasmic sites into two sets with the same number of sites as observed for the control region and the non-control region (p<0.001, Supplementary Figure S5). Thus, not only are more heteroplasmies shared in the control region than outside the control region, but control region heteroplasmies also have more similar MAFs.

**Figure 1:**
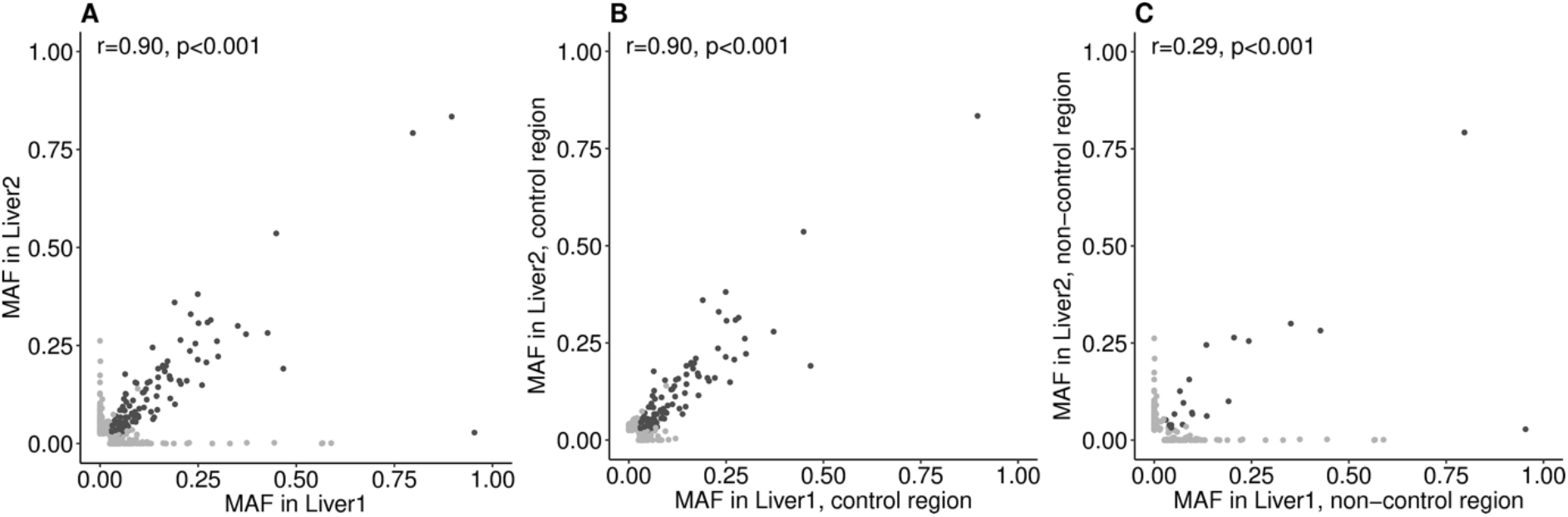
Correlation of MAFs at heteroplasmic sites in liver lobes. Each dot is one heteroplasmic site in one individual. Pearson’s correlation coefficient r is given. Heteroplasmic sites are compared in **A** liver sample 1 and 2, **B** the control region of liver sample 1 and 2, **C** the non-control region of liver sample 1 and 2. Heteroplasmic sites shared between liver sample 1 and 2 are in dark grey, non-shared sites in light grey.

**Table 1:** Heteroplasmic mutations in the control region and the non-control region and shared and non-shared heteroplasmies. p<0.001 (two-sided Fisher’s exact test).

| heteroplasms in liver1 and liver2 | control region | non-control region | total |
| --- | --- | --- | --- |
| shared | 115 | 21 | 136 |
| not shared | 95 | 260 | 355 |
| total | 210 | 281 |  |

### Potential mechanisms underlying the more frequent sharing in the control region

#### Frequent heteroplasmic sites

There are three sites that are frequently heteroplasmic in liver (Supplementary Table S1) and all are in the control region (sites 60, 72, and 94). To determine if these three sites are driving the higher correlation in MAF in the control region, we analyzed them separately. While these three sites did indeed show a high correlation in MAF between the two liver samples (Figure 2A, r = 0.91, p<0.001), the remaining sites in the control region still showed a significant correlation (Figure 2B, r=0.82, p<0.001) that is higher than the correlation for the non-control region (Figure 1C). To determine if the difference in correlations for the three sites vs. the remaining sites in the control region was statistically significant, we performed a subsampling test. As there were 113 occurrences of heteroplasmy at these three sites, we sampled 113 heteroplasmies at random from the control region, calculated the correlation in MAF between the two liver samples, and repeated this 1000 times. The r-value for the three sites was not significantly higher than the r-values for the random subsamples (Figure 2C), indicating that the higher correlation between MAF for the control region than for the rest of the genome is not driven solely by these three sites.

**Figure 2:**
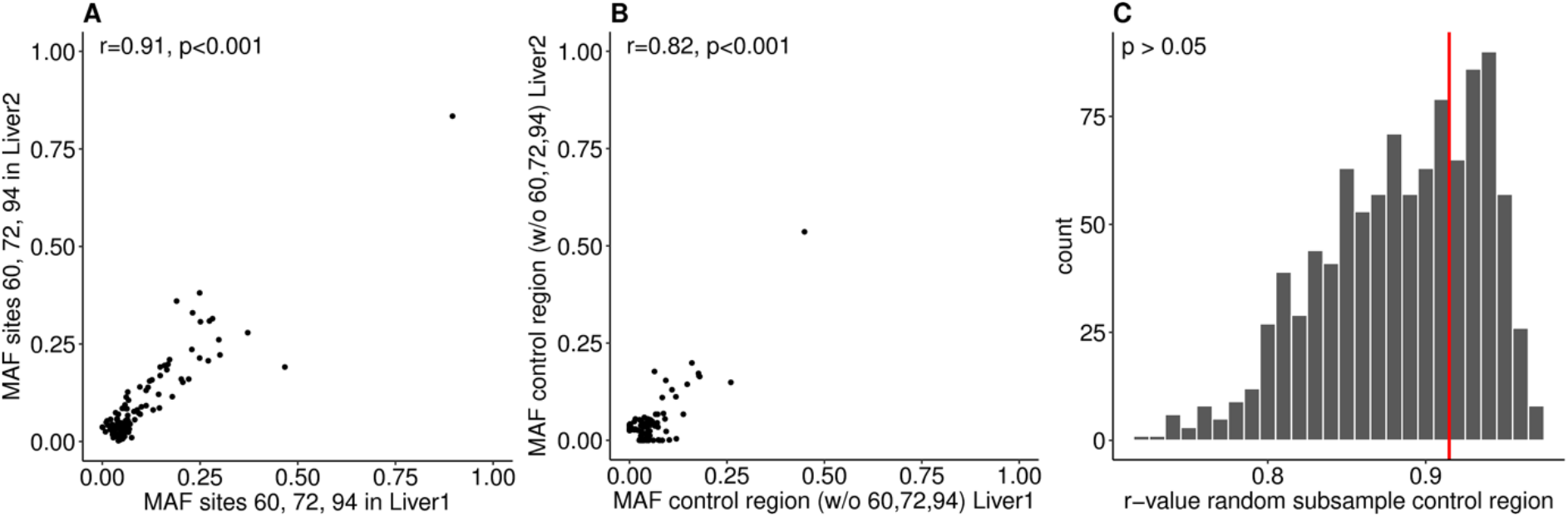
Heteroplasmy sharing in liver samples and correlation with single sites. MAFs per site in liver sample 1 and 2 for: **A** sites 60, 72 and 94; and **B** for all other heteroplasmic sites in the control region. Each dot is one heteroplasmic site in one individual. Pearson’s correlation coefficient r is given. **C** distribution of r-values for correlation of liver sample 1 and 2 MAFs per site for random subsamples of heteroplasmies in the control region (same number as the sum of sites 60, 72 and 94). The r-value for sites 60, 72 and 94 only is shown as a red bar.

#### 7S DNA

The control region includes the D-loop region, in which a third strand, the 7S DNA, displaces the heavy strand and binds to the light strand. Inferred heteroplasmies in the D-loop region might therefore reflect mutations in the 7S DNA rather than mutations in the mtDNA itself. To see if sequences from 7S DNA were likely to be present in the sequencing libraries, we estimated the relative mtDNA copy number from the capture-enrichment sequencing coverage of the D-loop region and the rest of the mtDNA molecule separately, as described before (Wachsmuth et al. 2016). As the 7S DNA has several starting and end points, we used the outer limits reported in the literature, namely from site 16,097 to site 191 (Roberti et al. 1998; Nicholls and Minczuk 2014). The D-loop did not exhibit a higher copy number than the other parts of the mtDNA genome, indicating that 7S DNA is unlikely to be present in the DNA libraries (p=0.30 and 0.41 for blood and liver, respectively, Supplementary Figure S6A). Furthermore, the correlation between the MAF for the two liver samples is almost as high for the D-loop region as it for the rest of the control region (Supplementary Figure S6 B,C: r=0.90 vs. r = 0.89). Hence, the significant correlation in MAFs in the control region is likely a phenomenon of the entire control region.

#### Higher mutation rate

We also tested if heteroplasmic sites showed a higher mutation rate than non-heteroplasmic sites by comparing the number of heteroplasmies to the inferred mutation rate at each site, based on observed polymorphism data (Soares et al. 2009). Heteroplasmic sites had significantly higher mutation rates than sites that were not heteroplasmic (p<0.001, Supplementary Figure S7) with the mutation rate being higher in the control region than outside the control region. However, mutation rates did not differ between shared vs. non-shared heteroplasmies (p>0.05, Supplementary Figure S7), suggesting that the mutation rate does not increase the probability of heteroplasmies to be shared.

#### Random genetic drift

If control region heteroplasmies are more likely to have arisen in a common progenitor cell of the cells sampled in the two liver lobes, then the higher correlation in MAF might reflect this common ancestry. We therefore tested if the correlation in MAF is compatible with expected amount of random genetic drift from a common progenitor cell by simulating mtDNA as a two-step model (Figure 3A) following (Jayaprakash et al. 2015). We started with different MAFs at the initial generation *n_0_* (5%, 10%, 20%, 50%) and different constant mitochondrial DNA copy numbers (1,000, 2,500, and 5,000) and sampled the difference in MAF 10, 20, 50, 100, and 250 generations after the initial split of the cells (Figure 3B). For all simulations, we observed a higher MAF difference with an increasing number of cell replications. This effect was more pronounced for higher MAF at generation *n_0_* and lower mtDNA copy number. Based on the observed mtDNA copy number (Figure 3C; mean=2,528) and MAF of shared heteroplasmies with the same consensus allele (Figure 3D; mean=10.8%), the most similar corresponding simulation is with a copy number of 2,500 and a MAF at generation *n_0_* of 10%; for this simulation, the variance in observed MAF differences was smaller than would be expected after 50 generations of random genetic drift (one-sided F-test for the equality of two variances, p<0.001). With an increasing number of cell replications, the simulated MAF difference further diverged from the observed values, which suggests that random genetic drift cannot explain the observed sharing.

**Figure 3:**
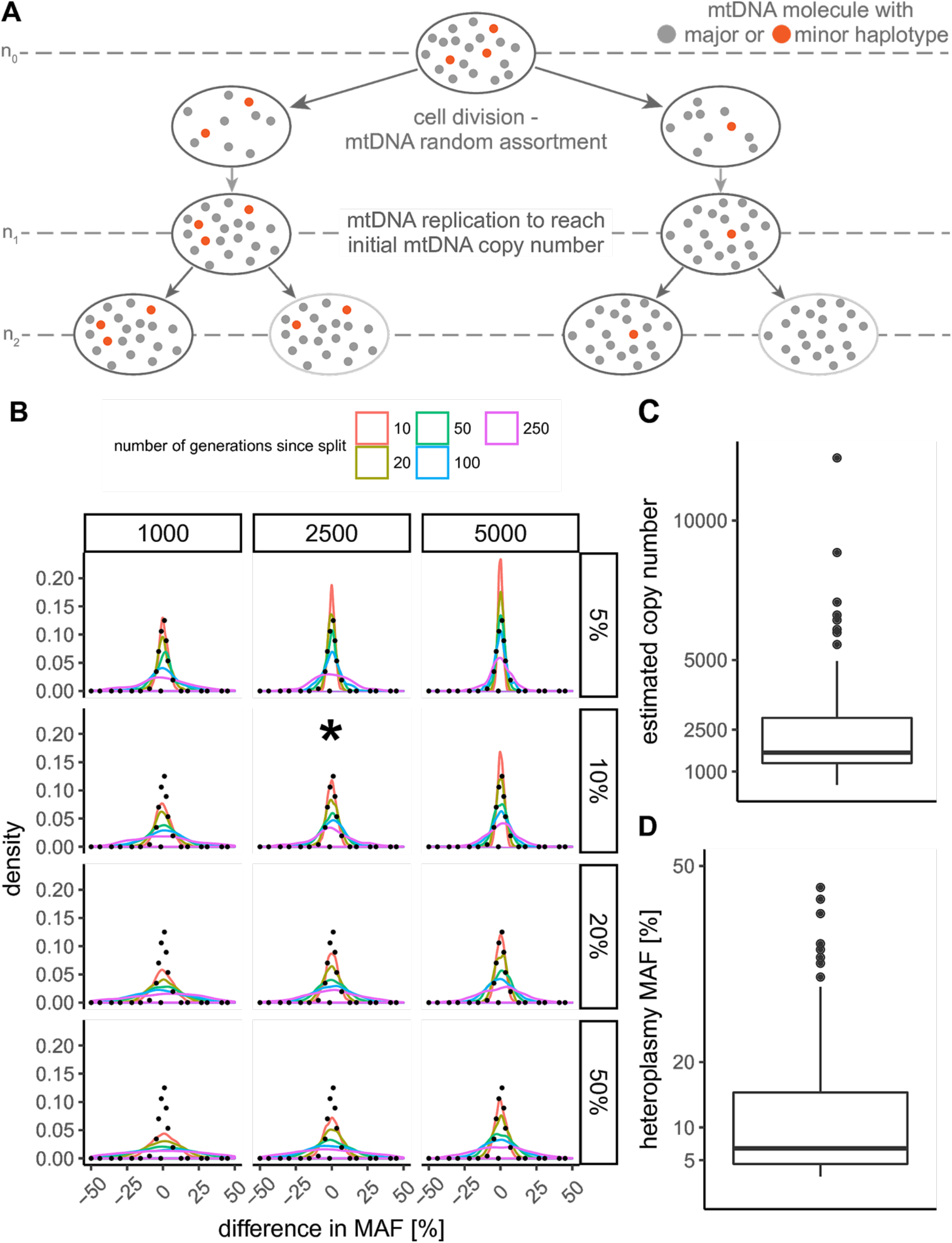
Expected difference in minor allele frequency (MAF) assuming random genetic drift. **A** schematic scheme of the two-step mtDNA replication model used for the simulations: first, all mtDNA molecules are segregated equally into two daughter cells followed by replication step to re-gain the same number of mtDNA copies as in the progenitor cell. After each generation *n_i_*, the MAF frequency of one random cell on each site of the pedigree is determined and the MAF difference between the two cells calculated. **B** expected difference in MAF after 10, 20, 50, 100, and 250 generations since the split of the two cells. Different mtDNA copy numbers (1,000, 2,500, and 5,000; kept constant throughout simulation) and initial MAFs (5%, 10%, 20%, and 50%) were used to simulate random genetic drift between two cells. For each combination of mtDNA copy number and initial MAF, 1000 replicates were simulated. The black, dotted line indicates the observed MAF difference between the shared liver heteroplasmies in the data set. The asterisk highlights the simulation with the parameters closest to the ones observed in the data set. **C** the mtDNA copy number distribution estimated from the capture-enriched sequencing data of the liver samples. The values were corrected using a correction “ratio” of 1/150 (Wachsmuth et al. 2016) to convert the relative to absolute mtDNA copy numbers. **D** the distribution of MAF in the liver heteroplasmies.

### Differences in MAF between corresponding liver samples are not correlated with age

We next investigated the influence of age on heteroplasmy sharing between different liver regions of an individual. Overall, the total number of different heteroplasmic sites in an individual increased with age for both the control region and the coding region (Supplementary Figure S8A, r=0.42 and adjusted p<0.001 both within and outside of the control region). However, the MAF at heteroplasmic sites did not increase with age (Supplementary Figure S8B, r=0.09 for the control region, −0.01 for outside of the control region, adjusted p>0.05). Although some heteroplasmies exhibit high MAFs only at ages above 50, many sites remain at low frequencies even at older ages (Supplementary Figure S8B). The hypothesis that random genetic drift has an effect on the difference in MAF of two corresponding liver samples would suggest that with increasing age the difference in MAF would increase, too. Yet, when testing for a correlation between the difference in MAF between two corresponding liver samples and age of the individual, we did not observe a significant correlation either within or outside the control region (Figure 4, r=0.06 within control region, r=−0.01 outside the control region, both p>0.05). In order to test whether, in contrast, specific sites have a significant correlation between the MAF difference and age, we separately calculated the correlation between MAF difference and age for the three most frequent heteroplasmy sites (sites 60, 72, and 94), for which at least 20 individuals were heteroplasmic (Supplementary Figure S9). The correlation between MAF and age was not significant for any of these three sites (adjusted p>0.05), which further supports the hypothesis that the difference in MAF does not increase with age, contrary to what would be expected from random genetic drift.

**Figure 4:**
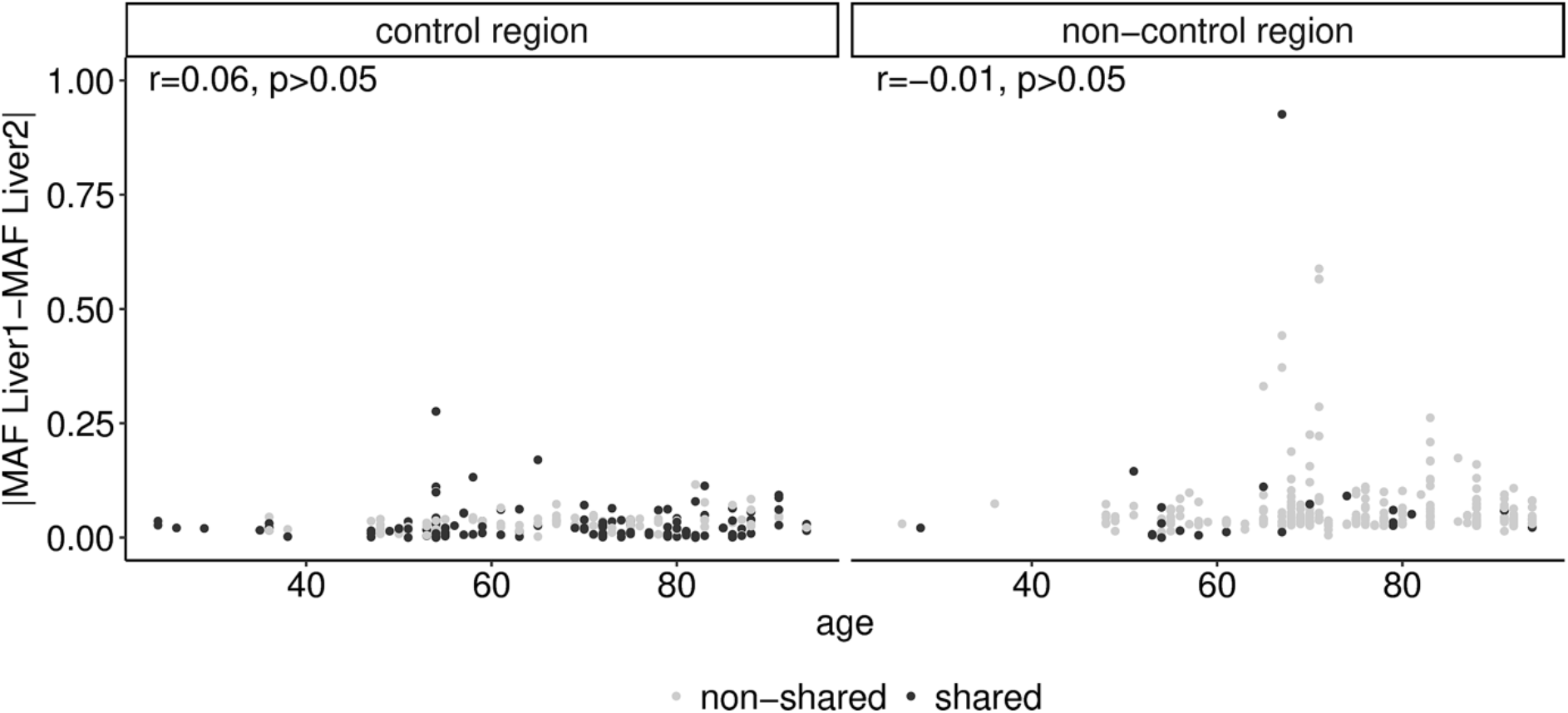
Minor allele frequency difference and age. The difference in minor allele frequency between liver sample 1 and 2 versus age plotted separately for sites within and outside the control region. Dark grey dots indicate sites that were shared (i.e. heteroplasmic in both liver samples from an individual), light grey dots indicate sites that were not shared.

### Synonymous heteroplasmies are more often shared than non-synonymous ones

Previous studies showed that liver has a significant excess of non-synonymous heteroplasmies which are predicted to have an impact on function (Li et al. 2015). We investigated this by calculating the ratio of non-synonymous heteroplasmies per non-synonymous site vs. synonymous heteroplasmies per synonymous site (hN/hS). In the absence of any selection this ratio has an expected value of 1; purifying selection results in values less than 1 and positive selection results in values greater than 1. As found previously (Li et al. 2015), the hN/hS ratio is significantly greater than 1 (adjusted p<0.05, Supplementary Figure S10A), indicating positive selection for non-synonymous heteroplasmies rather than relaxation of functional constraints (Li et al. 2015). Additionally, we observed a strong increase in the number of heteroplasmies occurring the coding region compared to the control region in individuals older than 60 years (Supplementary Figure S8A). We, therefore, investigated whether the excess in non-synonymous heteroplasmies is age-dependent (Supplementary Figure S10B). While both synonymous and non-synonymous heteroplasmies increased in number with increasing age, only non-synonymous heteroplasmies showed a significant difference between the age groups below 60 years and above 60 years (two-sided Mann-Whitney U test, adjusted p<0.05). We then asked if either synonymous or non-synonymous heteroplasmic sites were more likely to be shared between different liver samples. Although there were more than twice as many non-synonymous than synonymous heteroplasmies in the data set (163 vs. 62), there were significantly more synonymous sites shared than non-synonymous sites (11 vs. 6, Table 2). Accordingly, the median MAF difference between corresponding liver samples was on average higher for non-synonymous heteroplasmies than synonymous ones (median MAF difference of 4.4% to 3.8%), albeit not significantly so (p>0.05, Supplementary Figure S11A). These results suggest that non-synonymous heteroplasmies more often arise independently in different cells. However, neither non-synonymous nor synonymous heteroplasmies showed a significant correlation for the difference in MAF between corresponding liver samples with age (p>0.5, Supplementary Figure S11B).

**Table 2:** Non-synonymous and synonymous heteroplasmic mutations in the coding region and shared and non-shared heteroplasmies. p<0.001 (two-sided Fisher’s exact test)

| heteroplasmies in liver1 and liver2 | non-synonymous | synonymous | total |
| --- | --- | --- | --- |
| shared | 6 | 11 | 17 |
| not shared | 157 | 51 | 208 |
| total | 163 | 62 |  |

## DISCUSSION

### Heteroplasmies are shared across liver lobes

In this study, we compared mtDNA heteroplasmy patterns in samples collected from the two primary liver lobes of 83 individuals sampled at autopsy and found a significant correlation in heteroplasmy MAF between the two liver samples from an individual. Moreover, if sites are heteroplasmic in both liver samples of an individual, the MAFs at those sites tend to be similar independent of age. Previous studies found similar correlations in MAF between cell colonies derived from single cells and grown in culture (Raap et al. 2012; Jayaprakash et al. 2015), so our results extend these findings to different parts of the same organ within living individuals. In addition, we analyzed patterns of heteroplasmy sharing and MAF correlation across different parts of the mtDNA genome, which provide further information concerning the potential mechanism(s) behind these observations. In particular, we found significantly more sharing of heteroplasmic sites and higher MAF correlations for the control region than for the rest of the genome. We showed that these differences are not due to the presence of 7S DNA, nor are they driven by just a few sites. Moreover, within the coding region, we found significantly more sharing of heteroplasmies and higher MAF correlations at synonymous sites than at nonsynonymous sites.

These observations allow us to evaluate several potential explanations for this heteroplasmy sharing within liver. First, a trivial potential explanation is that a heteroplasmy could appear to be shared between liver lobes when it is actually present in blood. About 1 liter of blood per minute flows through the blood vessels of the liver (Wynne et al. 1989), so the DNA from the liver samples also contains DNA arising from blood cells. Heteroplasmies present in the blood cells could therefore be detected as heteroplasmies in the liver samples and hence seem to be shared across lobes. There are ~10^10^ liver cells in 10 g of liver tissue and 5 x 10^10^ blood cells in 10 ml of blood (Sender et al. 2016), and in liver autopsy samples there is approximately 10 ml blood per 100 g of tissue (Greenway and Stark 1971). The ratio of liver to blood cells in a liver autopsy sample is therefore about 1.8. As we additionally observed for our samples that the average mitochondrial copy number is five times higher in liver than in blood samples (Supplementary Figure S6a), the overall ratio of mitochondrial genomes from liver cells to those from blood cells is approximately 9 to 1. Therefore, a heteroplasmy in blood would need a MAF of at least 22.5% to be called a heteroplasmy in liver (minimum MAF = 2.5%) when it was actually totally absent in the liver samples. Only 8 out of the 541 heteroplasmies in blood have a MAF of ≥ 22.5%, of which four are shared across blood and both liver lobes (Supplementary Table S1). Moreover, the heteroplasmies that are commonly observed in liver (at positions 60, 72, and 94) are practically absent from blood; these results rule out experimental contamination with blood DNA as the primary driver for heteroplasmy sharing across liver lobes.

A second reason for heteroplasmy sharing could be a high mutational pressure for some sites, with de novo mutations at the same site occurring independently throughout the tissue. If this was the case, one would expect to see a higher correlation of MAFs for common heteroplasmic sites, as those would be under high mutational pressure. We showed that this is not the case for the most common heteroplasmic sites 60, 72, and 94 in the data set (Figure 2), and hence this explanation is unlikely. While we further observed a higher mutation rate for heteroplasmic sites than for non-heteroplasmic sites (Supplementary Figure S7), there was no difference in mutation rate between shared and non-shared heteroplasmies, suggesting that the mutation rate has little impact on heteroplasmy sharing.

Third, it has been shown that colonic stem cells with non-synonymous mtDNA mutations can expand clonally from a few cells and spread throughout a tissue by crypt fission (Greaves et al. 2006). During this process they retain the same level of mutant DNA on a cellular level. While a similar mechanism in liver could explain the results presented here, this process would require a cellular turnover on a huge scale throughout the entire liver, starting from just a few stem cells. This does not seem very likely, as clonal expansion was shown for single intestinal crypts only and is supposed to be rather slow (Greaves et al. 2006). Moreover, it is not clear to what extent liver regeneration is driven by stem cells vs. mature hepatocytes (Grompe 2014).

Fourth, heteroplasmy sharing could also derive from a pre-existing, inherited heteroplasmy (Guo et al. 2013; Payne et al. 2013; Rebolledo-Jaramillo et al. 2014; Li et al. 2015) that remains at similar frequencies across the tissue because drift (random changes in MAF) is limited. We tested this possibility, using a simulation scheme from a previous study (Jayaprakash et al. 2015), with different initial heteroplasmy frequencies, and mtDNA copy numbers that were kept constant (Berk and Clayton 1974). As expected, we observed a larger variance in MAF between daughter cells with increasing starting MAF and decreasing mtDNA copy number (Figure 3B). For the simulation results closest to our observed average mtDNA copy number and average heteroplasmy MAF (2,500 and 10%, Figure 3C-D), the observed difference in MAF between shared heteroplasmies was smaller than would be expected after random genetic drift acting on 50 mtDNA replication steps. However, more than these 50 mtDNA replication steps are expected to have taken place during the life of an average individual in our study. Assuming 3.61×10^11^ liver cells in an adult liver (Bianconi et al. 2013), a total of 39 hepatocyte cell replications (including mtDNA replication) are needed to obtain a full-size, adult liver from a single hepatocyte cell. After the development stage, the post-mitotic cells continue to replicate their mtDNA independently of cell replication (“relaxed” replication (Poovathingal et al. 2009)). The estimated half-life of mtDNA in post-mitotic cells ranges between 2-10 days (Miwa et al. 2008) and 30-300 days (Poovathingal et al. 2012). Assuming an mtDNA half-life of 30 days, there would be complete replacement of all mtDNA molecules of a cell within a year, given a mtDNA copy number of 2,500 per cell, so approximately one “relaxed” replication cycle occurs within the liver of an adult for every year of age. Thus, for the age range in our data set of 24 to 94 years, the liver samples would have gone through about 62 - 132 cell replications prior to sampling, and so the observed difference in MAF in the liver samples is significantly smaller than expected. Moreover, liver samples accumulated significantly more heteroplasmies with age (Supplementary Figure S8A), further arguing that shared heteroplasmies do not reflect pre-existing, inherited heteroplasmies. Also, there was no significant correlation of MAF difference with age, as would be expected with random genetic drift. In sum, our results extend to liver tissues the previous observations (Raap et al. 2012; Jayaprakash et al. 2015) that random genetic drift alone cannot explain heteroplasmy sharing between cells in culture.

Finally, an equilibrium of heteroplasmy across an entire tissue could be explained by an exchange of genetic material from mitochondria between cells. Cells can donate whole mitochondria to adjacent cells through nanotubes, but this has been suggested for distances up to 100 μM only and the exchange is often triggered by functional impairments in the acceptor cell (Rogers and Bhattacharya 2013). An additional way for cells to exchange DNA material could be the uptake of extracellular DNA material that is either secreted by healthy cells or is present as the remains of apoptotic cells (van der Vaart and Pretorius 2008). While the uptake and integration of cell-free nuclear DNA material has been shown (Basak et al. 2016), it is unclear whether cells would also accept mitochondrial DNA. However, studies of heteroplasmy at the single cell level (reviewed in (Yao et al. 2015)) do suggest the possibility of transfer between cells. Experiments with cell culture mixes of fluorescently labelled cell lines suggested the exchange of mtDNA between co-cultured partner cell lines, although the specific mechanism, either transfer of mitochondrial organelles or transfer of free mtDNA, could not be identified (Jayaprakash et al. 2015). Overall, such intercellular DNA exchange, followed by incorporation of mtDNA fragments into the mtDNA of the recipient cells, could account for the significant correlation we observe in MAF between liver lobes.

However, other aspects of our data are incompatible with the hypothesis of intercellular DNA exchange. In particular, intercellular exchange cannot explain the significantly higher number of shared heteroplasmies and correlation in MAF for the control region vs. the rest of the genome, unless one postulates that mtDNA fragments arising from the control region are either exchanged or incorporated between cells more frequently than mtDNA fragments arising from the rest of the genome. But even then, intercellular exchange cannot explain the significantly higher number of shared heteroplasmies and correlation in MAF for synonymous vs. non-synonymous heteroplasmies, as both should be exchanged at the same rate between cells.

Instead, our data suggest that even if intercellular exchange is occurring, selection must be involved in the sharing of heteroplasmies and correlation in MAF between liver lobes. Several aspects of the data suggest that selection influences heteroplasmies. In the coding region of the mtDNA, we observed a significant excess of non-synonymous vs. synonymous heteroplasmies (Table 2), more so than can be explained by relaxation of functional constraints on non-synonymous mutations. Moreover, the number of non-synonymous heteroplasmies increased significantly in individuals above 60 years. Overall, these results strongly suggest positive selection for nonsynonymous heteroplasmies in liver, as found previously (Li et al. 2015), and possibly reflecting the hypothesis of the “survival of the slowest” (deGrey 1997), which postulates that mitochondria with reduced respiratory function due to increasing mutations suffer less degradation from the production of reactive oxygen species (ROS). Hence, mitochondria that lack these mutations suffer ROS-related damage and are removed from cells, thereby resulting in an increase in frequency in mtDNAs with non-synonymous mutations that decrease mitochondrial function.

However, our results indicate a more complex role for selection in the different patterns of heteroplasmy sharing and MAF correlation across different regions of the mtDNA genome, in keeping with evidence from other studies. In mice and humans, significantly more synonymous than non-synonymous heteroplasmies are transmitted to the next generation (Stewart et al. 2008; Rebolledo-Jaramillo et al. 2014; Floros et al. 2018), suggesting selection against non-synonymous heteroplasmies during transmission. The notable tissue-specificity and allele-specificity of particular heteroplasmic sites in the control region also suggests a role for positive selection on heteroplasmies during aging (Samuels et al. 2013; Li et al. 2015). The increasing evidence for both purifying and positive selection acting on heteroplasmic variants warrants further investigation, particularly into the potential health-related consequences.

## MATERIAL AND METHODS

### Tissue collection and DNA extraction

Blood and liver were sampled at autopsy from 94 individuals (57 males, 37 females, age range: 24-94, mean: 63, median: 63). Two samples were taken from each liver, one from the right lobe and one from the left lobe. DNA was extracted as previously described (Li et al. 2015). The collection of samples and the experimental procedures were approved by the Ethics Commissions of the Rheinische Friedrich Wilhelms University Medical Faculty (Lfd. Nr. 097/15) and the University of Leipzig Medical Faculty (Az. 305-15-24082015).

### Virological assays and histological investigation

Human immunodeficiency virus (HIV) RNA and Hepatitis C virus (HCV) RNA concentration in blood was determined by using the Abbott RealTime^®^ HIV-1 and HCV systems and the m2000sp/m2000rt instruments according to the instructions of the manufacturer. For detection of Hepatitis B virus (HBV) DNA the Abbott RealTi*m*e^®^ HBV system was used. The 95% limit of detection (LOD_95_) of the HIV, HCV and HBV assay was 40 copies/mL, 12 IU/ mL, and 10 IU/ mL, respectively. If inhibitory effects on enzymatic reactions were present (detected via co-amplification of control RNA or DNA sequences), blood samples were re-tested at dilutions of 1/5, 1/10, and 1/15 (13 samples (13%) for HIV, 91 (93%) for HCV, and 8 (7%) for HBV load). For dilution, a plasma donation from a blood donor negative for HIV, HCV, and HBV was used. Of the diluted samples tested for HIV, one sample was diluted 1/5, nine had to be diluted 1/10, and two had to be diluted 1/15, lowering the LOD95 of the assay to 200, 400, and 600, respectively. In one sample, inhibition could not be eliminated by sample dilution. Of the diluted samples tested for HCV, one, 78, and two samples were diluted 1/5, 1/10, and 1/15, respectively, lowering the LOD_95_ of the assay to 60, 120, 180 IU/ml, respectively. In 10 additional samples no result could be achieved, even after dilution. Of the diluted samples tested for HBV load, 3, 2, and 2 samples were diluted 1/5, 1/10, and 1/15, respectively, lowering the LOD_95_ of the assay to 50, 100, and 150 IU/mL, respectively. All other samples were tested without any dilution.

Fat content of the liver was determined by histological investigations and Sudan staining (Mulisch M 2015). Tissues with <10% hepatocytes including fat droplets were considered low, 10-30% were medium, 31-50% were high fat and >50% were considered adipohepatic.

### Illumina library preparation and sequencing

Double-barcoded DNA libraries for sequencing were prepared and capture-enriched for mtDNA as previously described (Li et al. 2015). DNA was sequenced on the Illumina HiSeq platform in rapid mode with 95 bp paired-end reads. Bases were called with FreeIbis (Renaud et al. 2013) and reads were subsequently trimmed and merged using leeHom (Renaud et al. 2014).

### Heteroplasmy detection

Heteroplasmy was detected according to the DREEP pipeline (Li et al. 2010; Li and Stoneking 2012). First, heteroplasmies were called if: the minor allele frequency (MAF) for the most frequent minor allele was at least 2.5% on both the forward and reverse strand; the sequencing depth was at least 500-fold at a candidate site; and there were at least 10 reads supporting the minor allele on each strand. Additionally, we required a minimum heteroplasmic quality score of 10 on each strand. In order to discriminate a true heteroplasmy from sequencing error, the DREEP pipeline compares the minor allele pattern of any inferred heteroplasmic site to a database that comprises the minor allele patterns at this site from all other individuals in the study. When the majority of the samples are from a single tissue like liver in this study (two liver samples and one blood sample per individual), DREEP is prone to considered commonly heteroplasmic sites as elevated sequencing error and therefore under-estimates the heteroplasmic quality score for these. Thus, we used the information about commonly heteroplasmic sites from (Li et al. 2015) to flag these for both blood and liver tissue in this study, respectively. A site was considered commonly heteroplasmic in a tissue when at least five individuals were heteroplasmic for it. Based on this criterion, we ignored the heteroplasmic quality for sites 60, 72, 94, 185, 189, 203, 11,126, 16,093, and 16,126 for liver and 12,705 for blood and considered a site heteroplasmic if all other criteria were fulfilled. The following regions were excluded for heteroplasmy analysis: 302-316, 513-526, 566-573, and 16,181-16,194. We confirmed that all heteroplasmies were within a coverage between 20 % and 200 % of the average coverage of the sample. In addition, all samples that could have been contaminated with other samples during library preparation/extraction were removed. To detect such contamination, pairwise comparisons of all liver samples with each other as well as all blood samples with each other were performed. All three samples of an individual were removed if all of the following criteria were fulfilled for any pairwise comparison between individuals across all sites of the mitochondrial genome: 1) for at least 80% of the sites, for which two samples had different consensus alleles, the minor allele in the recipient sample was identical to the major allele in the donor sample; 2) the average MAF across these sites was at least 1%; and 3) at least 60% of all sites in the recipient sample, for which a minor allele was observed, were identified as heteroplasmies by the DREEP pipeline (Li et al. 2010; Li and Stoneking 2012). Furthermore, an additional filter for potential contamination was applied, in which the heteroplasmic sites for each sample were checked to see if five or more sites could be explained by contamination from another haplogroup. In total, ten individuals were removed in these contamination filter steps. Finally, we removed a single individual because its samples had more than 2% of the MT genome below a sequencing depth of 500-fold, the cut-off for being considered a heteroplasmy, and thus would have had a higher false negative rate than the other samples. Overall, we retained 83 individuals for the subsequent analyses. Minor allele frequencies were calculated with respect to the major allele in blood.

### Correlation analysis

Statistical analysis was performed using R (https://www.R-project.org), with analyses performed for the entire mtDNA genome and separately for the coding region (577-16,023), the control region (16,024-576) and the D-loop region (16,097-191). For correlation of MAFs, we selected only sites that were identified as heteroplasmies and passed our quality filters in at least one of the tissues. We then compared the MAFs of these heteroplasmies to the MAF in the other tissues of an individual, even if the site was not detected as a heteroplasmy in the other tissues. Pearson correlation coefficients were calculated for correlations between MAFs among samples as well as for correlations with age assuming a two-sided alternative hypothesis; the significance of the correlation was tested by randomly permuting the data. Permutation tests were also used to assess the association between specific sites or regions and minor allele sharing between liver lobes; all permutations were carried out 1000 times. Whenever categorical data were compared (e.g. synonymous vs. non-synonymous sites), two-sided Mann-Whitney U tests were used to test for significant differences. Fisher’s exact test was used to test for an association of sharing of heteroplasmic sites between liver lobes for the control region vs. non-control region and for synonymous vs. non-synonymous heteroplasmies using a two-sided alternative hypothesis. P-values were adjusted for multiple testing using Benjamini-Hochberg correction (Benjamini and Hochberg 1995) and highlighted in the text.

### Coverage across the mitochondrial genome

We used per-site coverage determined by the *filter_and_summary.pl* script of the DREEP pipeline (Li and Stoneking 2012) for each sample and calculated the average coverage across the D-loop region and across the rest of the mtDNA genome.

### Non-synonymous heteroplasmies

The hN/hS ratio (Li et al. 2015) was calculated by calculating the Ka/Ks ratio using *KaKs_Calculator 2.0* (Wang et al. 2010) between the revised Cambridge Reference Sequence (rCRS; Andrews, 1999 doi:10.1038/13779) and a mock sequence created by introducing all minor alleles of heteroplasmies into the coding region of the rCRS. A significance test was performed by randomly introducing the same substitutions as observed for heteroplasmies at any site of the coding region of rCRS and calculating the hN/hS ratio in comparison to the non-altered rCRS.

The potential functional impact of non-synonymous heteroplasmies was analyzed by overlapping the position of the heteroplasmy and its minor allele with the MitImpact database (Castellana et al. 2015) and comparing the *MutationAssessor* (Reva et al. 2011) score.

## DATA ACCESS

The raw sequencing data have been deposited with the European Nucleotide Archive under accession number PRJEB27731. The scripts for heteroplasmy detection and filtering can be found at http://dmcrop.sourceforge.net/ and https://github.com/alexhbnr/liverlobes_heteroplasmy.

## ACKNOWLEDGEMENTS

We thank Anne Haubner, Antje Weihmann, Barbara Höber and Melanie Grabmüller for technical support. We thank Enrico Macholdt, Michael Dannemann, and Martin Petr for valuable discussion. The research was funded by the Max Planck Society

## AUTHOR CONTRIBUTIONS

MS, BM, ML, AH and MW designed the study. BM collected the samples. RS performed DNA isolation and sequencing library preparation, AMEH performed virological assays and BM performed histological analyses. ML and AH wrote pipeline and executed heteroplasmy detection, AH and MW performed statistical analyses. AH, MW, AMEH and MS wrote the manuscript. All authors read and approved the final manuscript.

